# Low-efficiency conversion of proliferative glia into induced neurons by Ascl1 in the postnatal mouse cerebral cortex *in vivo*

**DOI:** 10.1101/2022.04.13.488173

**Authors:** Chiara Galante, Nicolás Marichal, Carol Schuurmans, Benedikt Berninger, Sophie Péron

## Abstract

The proneural transcription factor Achaete-scute complex-like 1 (Ascl1) is a major regulator of neural progenitor fate, implicated both in neurogenesis and oligodendrogliogenesis. Ascl1 has been widely used to reprogram non-neuronal cells into induced neurons. *In vitro*, Ascl1 induces efficient reprogramming of proliferative astroglia from the early postnatal cerebral cortex into interneuron-like cells. Here, we examined whether Ascl1 can similarly induce neuronal reprogramming of glia undergoing proliferation in the postnatal mouse cerebral cortex *in vivo*. Toward this, we targeted cortical glia at the peak of proliferative expansion (i.e., postnatal day 5) by injecting a retrovirus encoding for Ascl1 into the mouse cerebral cortex. In sharp contrast to the very efficient reprogramming *in vitro*, Ascl1-transduced glial cells were converted into doublecortin-immunoreactive neurons only with low efficiency *in vivo*. Interfering with the phosphorylation of Ascl1 by mutation of six conserved proline-directed serine/threonine phosphorylation sites (Ascl1SA6) has been previously shown to increase its neurogenic activity in the early embryonic cerebral cortex. We therefore tested whether transduction of proliferative glia with a retrovirus encoding Ascl1SA6 improved their conversion into neurons. While *in vitro* glia-to-neuron conversion was markedly enhanced, *in vivo* reprogramming efficiency remained low. However, both wild-type and mutant Ascl1 reduced the relative number of cells expressing the astrocytic marker glial fibrillary acidic protein (GFAP) and increased the relative number of cells expressing the oligodendroglial marker Sox10 *in vivo*. Together, our results indicate that the enhanced neurogenic response of proliferative postnatal glia to Ascl1SA6 versus Ascl1 observed *in vitro* is not recapitulated *in vivo*.

## INTRODUCTION

The postnatal mammalian brain is largely devoid of persistent neurogenesis, except from specialized niches such as the subependymal zone of the lateral ventricle and the subgranular zone of the dentate gyrus (Denoth-Lippuner and Jessberger, 2021). In all other brain regions, neurons lost due to disease or injury cannot be replaced, resulting in irreversible circuit dysfunction and functional impairments. Harnessing the neurogenic potential of glia to produce new neurons by direct lineage reprogramming has emerged as an approach for potential repair of diseased circuits in non-neurogenic brain areas such as the cerebral cortex (Peron and Berninger, 2015).

The basic helix-loop-helix (bHLH) transcription factor Ascl1 directly transactivates target genes and thereby orchestrates multiple aspects of cortical development including cellular proliferation, cell cycle exit, and neural differentiation (Castro et al., 2011; Guillemot and Hassan, 2017). Notably, Ascl1 controls GABAergic neurogenesis by regulating expression of homeobox genes of the distal-less gene family (*Dlx* genes) in progenitors of the ventral telencephalon (Casarosa et al., 1999; Poitras et al., 2007). We previously demonstrated that expression of Ascl1 in mouse postnatal cortical astrocytes *in vitro* was sufficient to reprogram them into functional neurons endowed with GABAergic identity (Berninger et al., 2007; Heinrich et al., 2010). Remarkably, Ascl1 can also reprogram cultured cells of human origin, including fibroblasts and pericytes, into neurons *in vitro* (Karow et al., 2012; Chanda et al., 2014). This raises the question whether it can also induce a neurogenic fate *in vivo*. For instance, it remains unknown whether glia of the cortical parenchyma can be reprogrammed into neurons *in vivo* with similar efficiency as *in vitro* when forced to express Ascl1 during their proliferative expansion, which peaks around postnatal day 5 (Ge et al., 2012).

Ascl1 function is tightly regulated by post-translational modifications, including phosphorylation (Dennis et al., 2019), which ultimately affects cell fate decisions. Notably, increased RAS/ERK signaling diverts Ascl1 from its neurogenic role and promotes a proliferative glial program (Li et al., 2014). bHLH transcription factors share an evolutionarily conserved serine/threonine phosphorylation residue in the L-H2 junction of the bHLH domain (Quan et al., 2016), but also harbor unique phosphorylation sites outside of this domain (Guillemot and Hassan, 2017). Most notably, similar to other bHLH transcription factors (Oproescu et al., 2021), Ascl1 is regulated by proline-directed serine threonine kinases, such as ERK (Li et al., 2014). Remarkably, preventing phosphorylation-dependent regulation of Ascl1 activity by mutating all six serines of the conserved serine-proline (SP) phospho-sites to alanine, a mutation here referred to as Ascl1SA6, has been found to increase its neurogenic activity in the embryonic day (E) 12.5 cerebral cortex (Li et al., 2014). This finding led us to hypothesize that using the Ascl1SA6 mutant variant could promote glia-to-neuron conversion both *in vitro* and *in vivo*.

Consistent with this hypothesis, our results show that Ascl1SA6 is more efficient than Ascl1 in converting postnatal cortical glia into neurons *in vitro*. However, Ascl1 and Ascl1SA6 had only limited reprogramming efficiency *in vivo*. Instead, we observed a reduction in the relative number of transduced cells expressing the astrocytic marker GFAP and a concomitant increase in the relative number of cells expressing the oligodendroglial lineage marker Sox10. This data suggests that, irrespective of its phosphorylation state, Ascl1 may preferentially promote an oligodendrogliogenic fate in proliferative postnatal cortical glia *in vivo*.

## MATERIALS AND METHODS

### Cell Culture

Postnatal cortical astrocytes were isolated from cortices of C57Bl6/J mice between postnatal day 5-7 days (P5-7), which were obtained from the Translational Animal Research Center of the University Medical Center Mainz.P5-P7 astrocytes were cultured as previously described (Heinrich et al., 2011; Sharif et al., 2021). Briefly, after isolation, cells were expanded for 7-10 days in Astromedium: Dulbecco’s Modified Eagles Medium, Nutrient Mixture F12 (DMEM/F12, Gibco, 21331-020); 10% Fetal Bovine Serum (FBS, Invitrogen, 10270-106); 5% Horse Serum (Invitrogen, 16050-130); 1x Penicillin/Streptomycin (Invitrogen, 15140122); 1x L-GlutaMAX Supplement (Invitrogen, 35050-0380); 1x B27 Supplement (Invitrogen, 17504001); and supplemented with 10ng/µl Epidermal Growth Factor (EGF; Peprotech, AF-100-15) and 10 ng/µl basic-Fibroblast Growth Factor (FGF-2; Peprotech, 100-18B). Cells were incubated at 37°C in 5% CO_2_. When cells reached 70-80% confluency, cells were detached with 0.05% Trypsin EDTA (Life Technologies, 15400054) for 5 min at 37°C. Cells were subsequently seeded onto poly-D-lysine hydrobromide-coated (PDL; Sigma, P0899) glass coverslips (12mm, Menzel-Gläser, 631-0713) in 24-well plates at a density of 50000-80000 cells/well in 500 µl Astromedium supplemented with 10 ng/µl EGF and 10 ng/µl FGF-2.

### Plasmids and retroviruses

Moloney Murine Leukaemia Virus (MMLV)-based retroviral vectors (Heinrich et al., 2011) were used to express Ascl1 and Ascl1SA6 under control of the chicken β-actin promoter with a cytomegalovirus enhancer (pCAG). A GFP or DsRed reporter was cloned in behind an Internal Ribosome Entry Site (IRES). To generate the pCAG-Ascl1-IRES-DsRed/GFP and pCAG-Ascl1SA6-IRES-DsRed/GFP retroviral constructs, a cassette containing the coding sequences flanked by attL recombination sites was generated through the excision of the coding sequences for Ascl1 and Ascl1SA6 from the pCIG2 parental vectors (Li et al., 2014) via XhoI/SalI double restriction. Isolated fragments were inserted into the pENTRY1A Dual Selection (Invitrogen) intermediate vector linearized via SalI. The final retroviral constructs were subsequently obtained via recombination catalyzed by the LR Clonase II (Invitrogen, 11791020), which substituted the ccdB cassette in the destination vector pCAG-ccdB-IRES-DsRed or pCAG-ccdB-IRES-GFP with Ascl1 or Ascl1SA6 coding sequences. Transduction with MMLV-based retroviral vectors encoding only the fluorescent protein GFP or DsRed behind an IRES under control of pCAG promoter (pCAG-IRES-DsRed/pCAG-IRES-GFP) (Heinrich et al., 2011) was used for control experiments. Viral particles were produced using gpg helper free packaging cells to generate Vesicular Stomatitis Virus Glycoprotein (VSV-G)-pseudotyped retroviral particles (Ory et al., 1996). Viral stocks were titrated by transduction of HEK293 cultures. Viral titers used were in the range of 10^7^ TU/ml.

### Retroviral transduction

After seeding the cells and letting them attach for 4h in the incubator, cells were transduced with 1 µl retrovirus/well and incubated at 37°C in 8% CO_2._ One day later, treated medium was removed and substituted with 500 µl of B27 Differentiation Medium: DMEM/F12; 1x Penicillin/Streptomycin; 1x L-GlutaMAX Supplement; 1x B27 Supplement. Cells were treated again with 1 µl/well of retrovirus. One day later, the culture volume was brought to 1 ml/well with fresh B27 Differentiation Medium. Cells were kept in culture for a total of 7 days *in vitro* before fixation for immunocytochemical analyses.

### Immunocytochemistry

Cells were fixed with 4% paraformaldehyde (PFA, Sigma, P6148) for 10-15 min and washed 3 times with 1xPBS (Gibco, 70013-016) before storage at 4°C. Washed cells were first incubated for 1 h at room temperature (RT) with blocking solution (3% bovine serum albumin [BSA, Sigma, A7906] and 0.5% Triton X-100 [Sigma, X100] in 1xPBS) and then with primary antibodies for 2-3 h at RT. After 3 washes with 1xPBS, cells were incubated with secondary antibodies for 1h at RT. Cells were then counterstained with DAPI (Sigma, D8417) diluted 1:1000 in blocking solution, then washed 3 time in 1xPBS before being mounted with Aqua Polymount (Polysciences, 18606-20). The following primary antibodies were used: β-Tubulin III (Mouse IgG2b, 1:1000, Sigma, T8660); Green Fluorescent Protein (GFP, Chicken, 1:300, AvesLab, GFP-1020); GFAP (rabbit, 1:1000, Dako, Z0334); Red Fluorescent Protein (RFP, rat, 1:400, Chromoteck, 5F8). Secondary antibodies were diluted 1:1000 and were conjugated to: A488 anti-chicken (donkey, Jackson Immunoresearch, 703-545-155); Cy3 anti-mouse (goat, Dianova, 115-165-166); Cy3 anti-rat (goat, Dianova, 112-165-167); Cy5 anti-rabbit (goat, Dianova, 111-175-144).

### Animals and Animal Procedures

The study was performed in accordance with the guidelines of the German Animal Welfare Act and the European Directive 2010/63/EU for the protection of animals used for scientific purposes and was approved by the Rhineland-Palatinate State Authority (permit number 23 177 07-G15-1-031). For retroviral injections, male and female C57Bl6/J pups kept with their mother were purchased from Janvier Labs. Mice were kept in a 12:12 h light-dark cycle in Polycarbonate Type II cages (350 cm^2^). Animals were provided with food and water *ad libitum* and all efforts were made to reduce the number of animals and their suffering. Before the surgery, animals received a subcutaneous injection of Carprofen (Rimadyl®, Zoetis, 4 mg/kg of body weight, in 0.9% NaCl [Amresco]). Anaesthesia was induced by intraperitoneal (i.p.) injection of a solution of 0.5 mg/kg Medetomidin (Pfizer), 5 mg/kg Midazolam (Hameln) and 0.025 mg/kg Fentanyl (Albrecht) in 0.9% NaCl. Viruses were injected in the cerebral cortex using glass capillaries (Hirschmann, 9600105) pulled to obtain a 20 µm tip diameter. Briefly, a small incision was made on the skin with a surgical blade and the skull was carefully opened with a needle. Each pup received a volume of 0.5-1 μl of retroviral suspension targeted to the somatosensory and visual cortical areas. After injection, the wound was closed with surgical glue (3M Vetbond, NC0304169) and anesthesia was terminated by i.p. injection of a solution of 2.5 mg/kg Atipamezol (Pfizer), 0.5 mg/kg Flumazenil (Hameln) and 0.1 mg/Kg Buprenorphin (RB Pharmaceutials) in 0.9% NaCl. Pups were left to recover on a warm plate (37°C) before returning them to their mother. Recovery state was checked daily for a week after the surgery.

### Tissue preparation and immunohistochemistry

Animals were lethally anesthetized with a solution of 120 mg/kg Ketamine (Zoetis) and 16 mg/kg Xylazine (Bayer) (in 0.9% NaCl, i.p.) and transcardiacally perfused with pre-warmed 0.9% NaCl followed by ice-cold 4% paraformaldehyde (PFA, Sigma, P6148). The brains were harvested and post-fixed for 2 h to overnight in 4% PFA at 4°C. Then, 40 μm thick coronal sections were prepared using a vibratome (Microm HM650V, Thermo Scientific) and stored at -20°C in a cryoprotective solution (20% glucose [Sigma, G8270], 40% ethylene glycol [Sigma, 324558], 0.025% sodium azide [Sigma, S2202], in 0.5 X phosphate buffer [15mM Na_2_HPO_4_·12H_2_O [Merck, 10039-32-4]; 16mM NaH_2_PO_4_ ·2H_2_O [Merck, 13472-35-0]; pH 7.4]).

For immunohistochemistry, brain sections were washed three times for 15 min with 1X TBS (50mM Tris [Invitrogen, 15504-020]; 150 mM NaCl [Amresco, 0241]; pH7.6) and then incubated for 1.5 h in blocking solution: 5% Donkey Serum (Sigma, S30); 0.3% Triton X-100; 1X TBS. Slices were then incubated with primary antibodies diluted in blocking solution for 2-3 h at RT, followed by an overnight incubation at 4°C. After three washing steps with 1X TBS, slices were incubated with secondary antibodies diluted blocking solution for 1 h at RT. Slices were washed twice with 1X TBS, incubated with DAPI dissolved in 1X TBS for 5 min at RT and washed three times with 1X TBS. For mounting, slices were washed two times with 1X Phosphate Buffer (30 mM Na_2_HPO_4_·12H_2_O [Merck, 10039-32-4]; 33 mM NaH_2_PO_4_ ·2H_2_O [Merck, 13472-35-0]; pH 7.4) and were dried on Superfrost (Thermo Fisher Scientific) microscope slides. Sections were further dehydrated with toluene and covered with cover-glasses mounted with DPX mountant for histology (Sigma, 06522) or directly mounted with Prolong™Gold (Invitrogen, P36930). The following primary antibodies were used: Achaete scute-like1 (Ascl1, mouse IgG1, 1:400, BD Pharmingen, 556604); Doublecortin (DCX, goat, 1:250, Santa Cruz Biotechnology, sc-8066); Green Fluorescent Protein (GFP, chicken, 1:1000, AvesLab, GFP-1020); Glial Fibrillary Acidic Protein (GFAP, rabbit, 1:300, Dako, Z0334); Ionized calcium-binding adapter molecule 1 (Iba1, rabbit, 1:800, Wako, 16A11); mCherry (chicken, 1:300, EnCor Biotechnology, CPCA-mCherry); Red Fluorescent Protein (RFP, rabbit, 1:500, Biomol, 600401379S); SRY-Box 10 (Sox10, goat, 1:100, Santa Cruz Biotechnology, sc-17342). Secondary antibodies were made in donkey and conjugated with: A488 (anti-chicken, 1:200, Jackson Immunoresearch, 703-545-155); A488 (anti-rabbit, 1:200, Invitrogen, A21206); A647 (anti-rabbit, 1:500, Invitrogen, A31573); A488 (anti-mouse, 1:200, Invitrogen, A21202); A647 (anti-mouse, 1:500, Invitrogen, A31571); Cy3 (anti-chicken, 1:500, Dianova, 703-165-155); Cy3 (anti-goat, 1:500, Dianova, 705-165-147); Cy3 (anti-mouse, 1:500, Invitrogen, A10037); Cy3 (anti-rabbit, 1:500, Dianova, 711-165-152); Cy5 (anti-goat, 1:500, Dianova, 705-175-147).

### Imaging and data analysis

Images were acquired using a TCS SP5 (Leica) confocal microscope (Institute of Molecular Biology, Mainz, Germany) equipped with four PMTs, four lasers (405 Diode, Argon, HeNe 543, HeNe 633) and a fast-resonant scanner. Images were taken with a 20x dry objective (NA 0.7) or a 40x oil objective (NA 1.3). For imaging of brain sections, serial Z-stacks spaced at 0.3 μm-1.25 μm distance were acquired to image the whole thickness of the section. Alternatively, imaging was performed using an Axio Imager.M2 fluorescent microscope equipped with an ApoTome (Zeiss) at a 20x dry objective (NA 0.7) or a 63x oil objective (NA 1.25).

For *in vitro* experiments, biological replicates (n) were obtained from independent cultures prepared from different animals. For each n, the value corresponds to the mean value of two technical replicates (i.e., two coverslips). Cell quantifications were performed on 4×4 tile scans (individual tile size: 624,70×501,22 μm). For *in vivo* experiments, n corresponds to the number of animals. Quantifications were performed on equally spaced sections (240 or 480 µm) covering the whole area with transduced cells. Tile scans were acquired with a serial Z-stack spaced at 1.25 μm distance.

For images used for illustration, the color balance of each channel was uniformly adjusted in Photoshop (Adobe). If necessary, Lookup Tables were changed to maintain uniformity of color coding within figures. When appropriate, a median filter (despeckle) was applied in Fiji to pictures presenting salt-and-pepper noise, and noise was filtered via removal of outlier pixels.

Multiple sequence alignment of the Ascl1 and Ascl1SA6 protein sequences was performed in Clustal Omega (RRID:SCR_001591).

### Statistical analysis

The number of independent experiments (n) and number of cells analyzed are reported in the main text or figure legends. Data are represented as means ± SD. Statistical analysis was performed in SPSS Statistics 23 V5 (IBM). Normality of distribution was assessed using Shapiro-Wilk test and the significance of the differences between groups was analyzed by One-Way ANOVA followed by Bonferroni post-hoc test. P-values are indicated in the figures or figure legends. Graphs were prepared in GraphPad Prism 5.

## RESULTS

### Ascl1SA6 improves neuronal reprogramming efficiency from cultured postnatal cortical astroglia compared to wild-type Ascl1

Our earlier work showed that Ascl1 can reprogram cultured postnatal astroglia into neurons (Berninger et al., 2007; Heinrich et al., 2010). More recently, overexpression of Ascl1SA6 in embryonic cortical progenitors was found to enhance neuronal differentiation compared with wild-type Ascl1 (Li et al., 2014). Here, we examined whether Ascl1SA6 can increase the glia-to-neuron reprogramming capacity of Ascl1 in postnatal astroglial cultures. For this purpose, we first cloned murine Ascl1 and Ascl1SA6 sequences into retroviral vectors for transduction of proliferative glia. Figure 1A depicts the six serine-to-alanine (SA6) substitutions resulting from the targeted mutation of the Ascl1 coding sequence. Astroglial cultures prepared from P5 mice were transduced with retroviruses encoding for Ascl1 or Ascl1SA6 together with a reporter gene (GFP or DsRed). A retrovirus encoding only a reporter gene was used as control (Figure 1B). Neuronal reprogramming efficiency was evaluated seven days after transduction by immunocytochemistry directed against the neuronal marker β-Tubulin III. After transduction with control virus, virtually no β-Tubulin III-immunoreactive cells were found (0.1±0.2%, 1398 transduced cells analysed, n=3 biological replicates; Figure 1C,D). In contrast, consistent with our previous findings (Berninger et al., 2007; Heinrich et al., 2010), astrocytes transduced with Ascl1 acquired a neuronal-like elongated morphology and expressed β-Tubulin III (27.3±3.8%, 3061 transduced cells analysed, n=4 biological replicates; Figure 1C,D). Strikingly, the proportion of converted cells doubled upon overexpression of Ascl1SA6 (51.1±7.0%, 3462 transduced cells analysed, n=3 biological replicates; Figure 1C,D).

**Figure 1.**
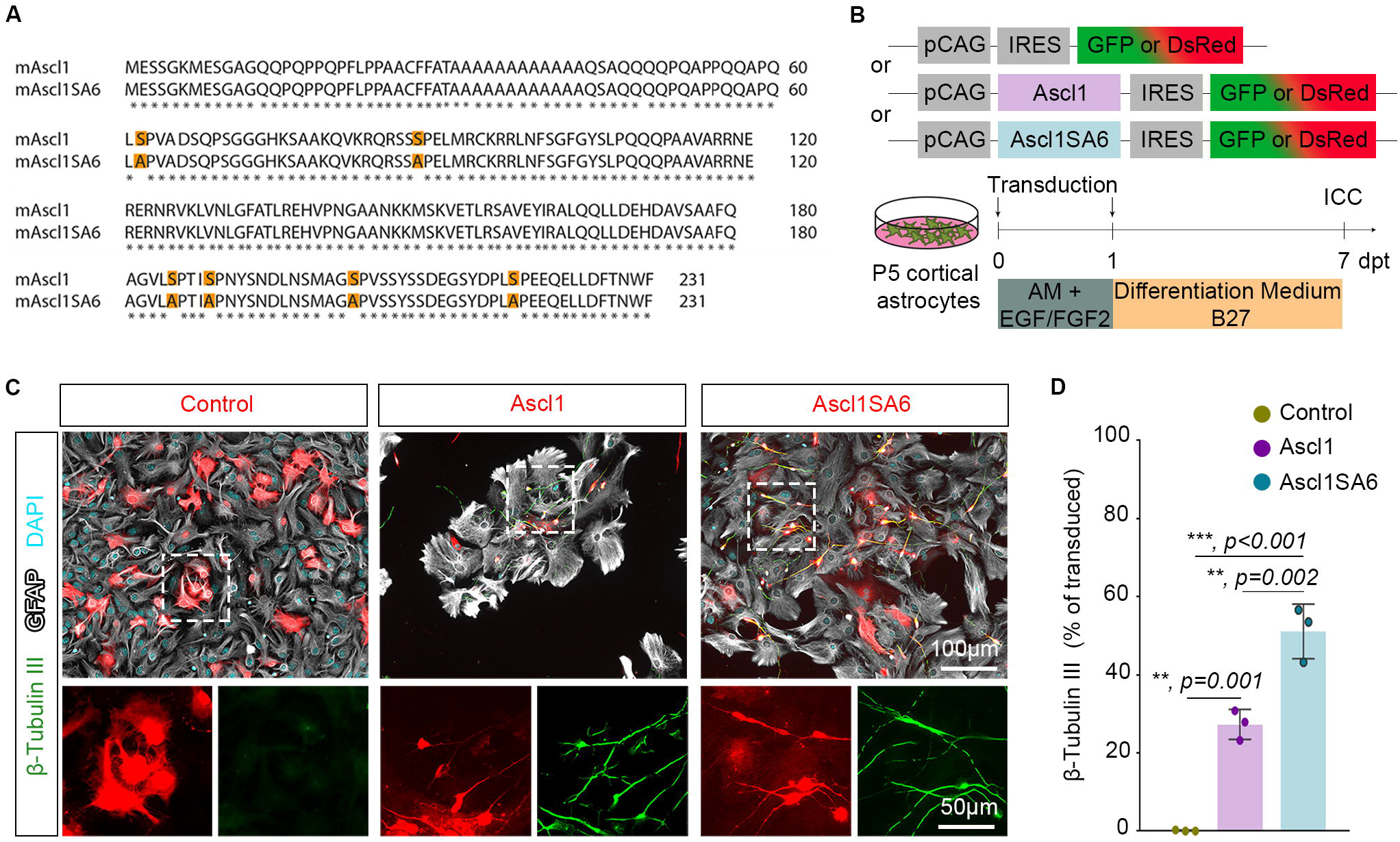
Ascl1SA6 induces more efficient glia-to-neuron reprogramming *in vitro*. **(A)** Multiple sequence alignment depicts the 6 serine residues mutated in the sequence of mouse Ascl1 (mAscl1) to generate the mutant Ascl1SA6. **(B)** Experimental scheme. Postnatal day 5 (P5) cortical astrocytes cultures were transduced with pCAG-Ascl1-IRES-DsRed/GFP or pCAG-Ascl1SA6-IRES-DsRed/GFP retroviral constructs, or pCAG-IRES-DsRed/pCAG-IRES-GFP as a control. Seven days later, cells were fixed for immunocytochemical (ICC) analysis. **(C)** Representative pictures of the cultures transduced with control, Ascl1 or Ascl1SA6-encoding retroviruses. In control, transduced cells (in red) exhibit an astroglia-like morphology, express GFAP (in grey) and lack β-Tubulin III (in green) expression. Conversely, Ascl1- and Ascl1SA6-transduced cells develop neuronal morphological hallmarks and acquire β-Tubulin III expression. **(D)** Quantification of the percentage of transduced cells expressing β-Tubulin III indicates higher reprogramming efficiency with Ascl1SA6. AM, Astromedium; dpt, days post-transduction; ICC, immunocytochemistry; P, postnatal day.

These results indicate an increased neurogenic potential of Ascl1SA6 in glia-to-neuron conversion when expressed in postnatal astrocytes *in vitro*.

### Ascl1 or Ascl1SA6 converts postnatal cortical glia into neurons with low efficiency *in vivo*

We next tested whether, like *in vitro*, proliferative cortical glia can be efficiently reprogrammed towards a neuronal fate *in vivo*. Cortical glia greatly expands during the first postnatal week by local proliferation (Ge et al., 2012; Clavreul et al., 2019), To target proliferative glia, retroviruses were injected into the cerebral cortex at postnatal day five (P5). We then analyzed the identity of the transduced cells by immunohistochemical analysis at three days post injection (3 dpi), first with the control virus only (Figure 2A). We found that virtually all transduced cells were immunopositive for glial markers (Figure 2B). The majority of transduced cells were immunoreactive for the astroglial marker GFAP (62.8±8.1%, 753 transduced cells analysed, n=3 mice; Figure 2B,C), and the remaining were oligodendroglial cells immunoreactive for Sox10 (32.3±6.1%, 753 transduced cells analysed, n=3 mice; Figure 2B,C). Rarely, we found transduced cells immunoreactive for the microglial marker Iba1 (1.0±0.9%, 578 transduced cells analysed, n=3 mice; Figure 2B,D). Importantly, none of the control-transduced cell expressed the immature neuronal marker DCX (0.0±0.0%, 578 transduced cells analysed, n=3 mice; Figure 2B,D). These results indicate that retroviruses injected in the P5 mouse cerebral cortex *in vivo* specifically transduce astroglial and oligodendroglial lineage cells.

**Figure 2.**
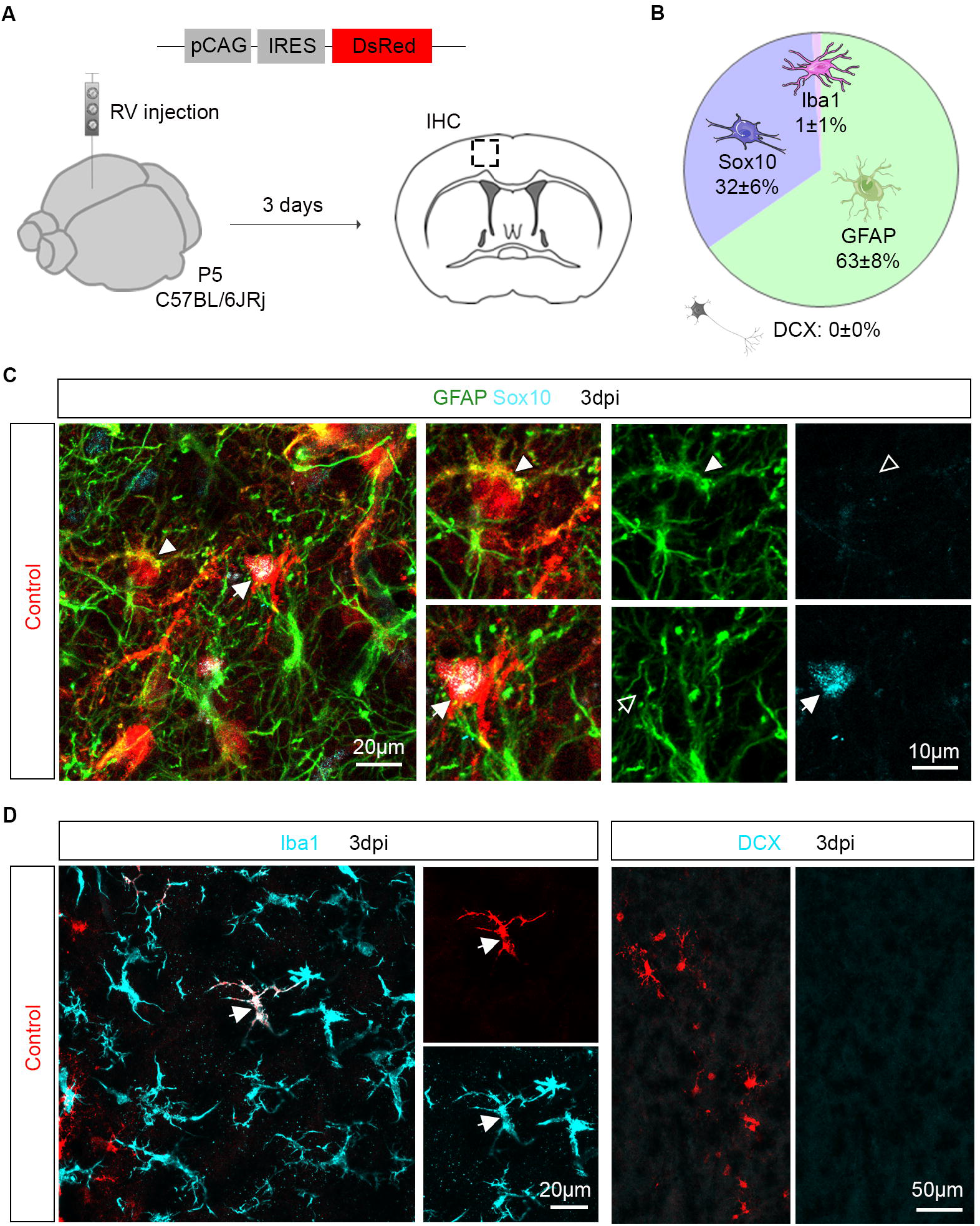
Retroviruses injected in the postnatal cerebral cortex specifically transduce glial cells. **(A)** Experimental scheme. A control retrovirus pCAG-IRES-DsRed was injected in the cerebral cortex of P5 mice and immunohistochemical analysis was performed 3 days later. **(B)** Pie chart showing the relative number of oligodendroglial (Sox10-positive), astroglial (GFAP-positive), microglial (Iba1-positive) and neuronal (DCX-positive) cells among transduced cells. **(C)** Confocal images depicting control-transduced cells (in red) co-expressing GFAP (in green, arrowheads, upper insets) or Sox10 (in blue, arrows, lower insets). **(D)** Confocal images depicting control-transduced cell (in red) co-expressing Iba1 (in cyan, left panel). No control-transduced cells expressing DCX were found (in cyan, right panel). Full arrows/arrowheads indicate marker-positive cells; empty arrows/arrowheads indicate marker-negative cells. IHC, immunohistochemistry; RV, retrovirus.

We next injected control, Ascl1- or Ascl1SA6-encoding retroviruses and investigated whether these bHLH genes could reprogram P5 proliferative glia into neurons by immunohistochemical analysis at 12 dpi. Ascl1 was effectively expressed in cells transduced with Ascl1 and Ascl1SA6, while absent from control-transduced cells (Figure 3A). Control-transduced cells lacked DCX expression (0.0±0.0%, 2157 transduced cells analysed, n=3 mice; Figure 3B,C), confirming that the control vector did not induce a cell fate switch. In contrast to our findings *in vitro*, Ascl1- and Ascl1SA6-transduced cells largely remained immunonegative for DCX (Figure 3B), with only a small number of cells exhibiting an immature neuron-like morphology and expressing DCX (Ascl1: 4.6±1.6%, 720 transduced cells analysed, n=3 mice, and Ascl1SA6: 6.9±0.2%, 409 transduced cells analysed, n=3 mice) (Figure 3B,C). Together, our results indicate that despite the strong neurogenic potential of Ascl1 and Ascl1SA6 *in vitro* (Figure 2), these bHLH genes can only reprogram postnatal cortical glia into neurons with low efficiency *in vivo*.

**Figure 3.**
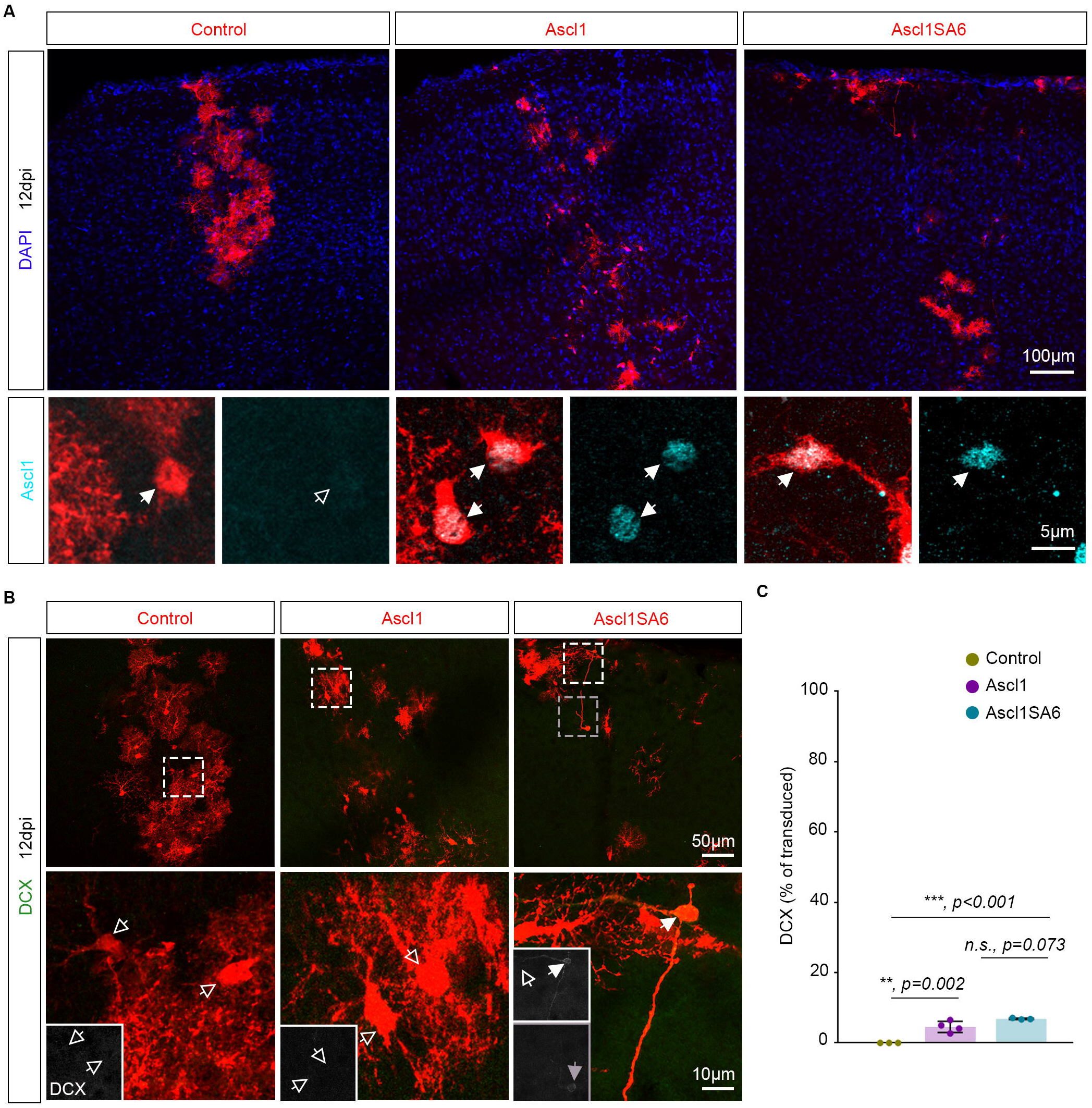
Ascl1 and Ascl1SA6 convert postnatal glia into neurons with low efficiency *in vivo*. **(A)** Immunohistochemistry confirmed the absence of Ascl1 in control-transduced postnatal cortical glia and efficient Ascl1 induction by pCAG-Ascl1-IRES-DsRed/GFP and pCAG-Ascl1SA6-IRES-DsRed/GFP retroviruses. **(B)** Confocal images depicting the maintenance of a glial morphology and lack of DCX induction in most transduced cells with only a few transduced cells acquiring a neuronal morphology and expressing DCX following transduction with Ascl1 and Ascl1SA6. **(C)** Quantification of the percentage of transduced cells expressing DCX at 12dpi indicates that Ascl1 and Ascl1SA6 induce neurogenesis from postnatal cortical glia with low efficiency. Full arrows/arrowheads indicate marker-positive cells; empty arrows/arrowheads indicate marker-negative cells. Dpi, days post infection.

### Ascl1 expression in postnatal cortical glia increases the relative number of cells expressing oligodendroglial markers

Given that only a few Ascl1 or Ascl1SA6 transduced cells were converted into neurons, we examined whether the remaining cells nevertheless had responded to the reprogramming factors and downregulated glial markers. We therefore analyzed the expression of the pan-oligodendroglial marker Sox10 and the astroglial marker GFAP in Ascl1- and Ascl1SA6-transduced cells (Figure 4A-C). Consistent with our analysis at 3 dpi (Figure 2), control-transduced cells at 12 dpi were glial cells, predominantly astrocytes, with two thirds of the cells expressing GFAP (63.3 ± 11.3%, 1885 transduced cells analysed, n=3 mice) and another third expressing Sox10 (35.6±12.1%, 1885 transduced cells analysed, n=3 mice; Figure 4A,C). As expected, the expression of GFAP and Sox10 was mutually exclusive in control-transduced cells (0.3±0.6% of GFAP/Sox10-positive cells, 1885 transduced cells analysed, n=3 mice; Figure 4B,C and Supplementary Movie 1). Following transduction with both Ascl1 variants, we observed a marked alteration in the expression of glial markers. Strikingly, only one fifth of transduced cells expressed exclusively GFAP (Ascl1, 18.7±3.1%, 848 transduced cells analysed, n=4 mice; Ascl1SA6, 20.4±6.1%, 573 transduced cells analysed, n=3 mice). Interestingly, in Ascl1-transduced cells, the reduction in GFAP expression was concomitant with a large increase in the relative number of Sox10-only expressing cells (70.0±7.7%, 848 transduced cells analysed, n=4 mice; Figure 4C). The same trend was observed following Ascl1SA6 overexpression (50.7±3.1%, 573 transduced cells analysed, n=3 mice; Figure 4C), albeit without reaching statistical significance. Instead, a significant increase in the relative number of cells co-expressing Sox10 and GFAP was observed in Ascl1SA6-transduced cells (Figure 4B,C and Supplementary Movie 3). The detection of GFAP/Sox10-immunopositive cells following transduction with both Ascl1 variants (Ascl1, 4.5±2.6%, 848 transduced cells analysed, n=4 mice; Ascl1SA6, 17.1±6.0%, 573 transduced cells analysed, n=3 mice, Figure 4B,C and Supplementary Movies 2 and 3) may capture cells in a “mixed” glial state. These results indicate that although largely failing to redirect towards neurogenesis, proliferative glial cells appear to be responsive to Ascl1 or Ascl1SA6 by turning on Sox10 expression.

**Figure 4.**
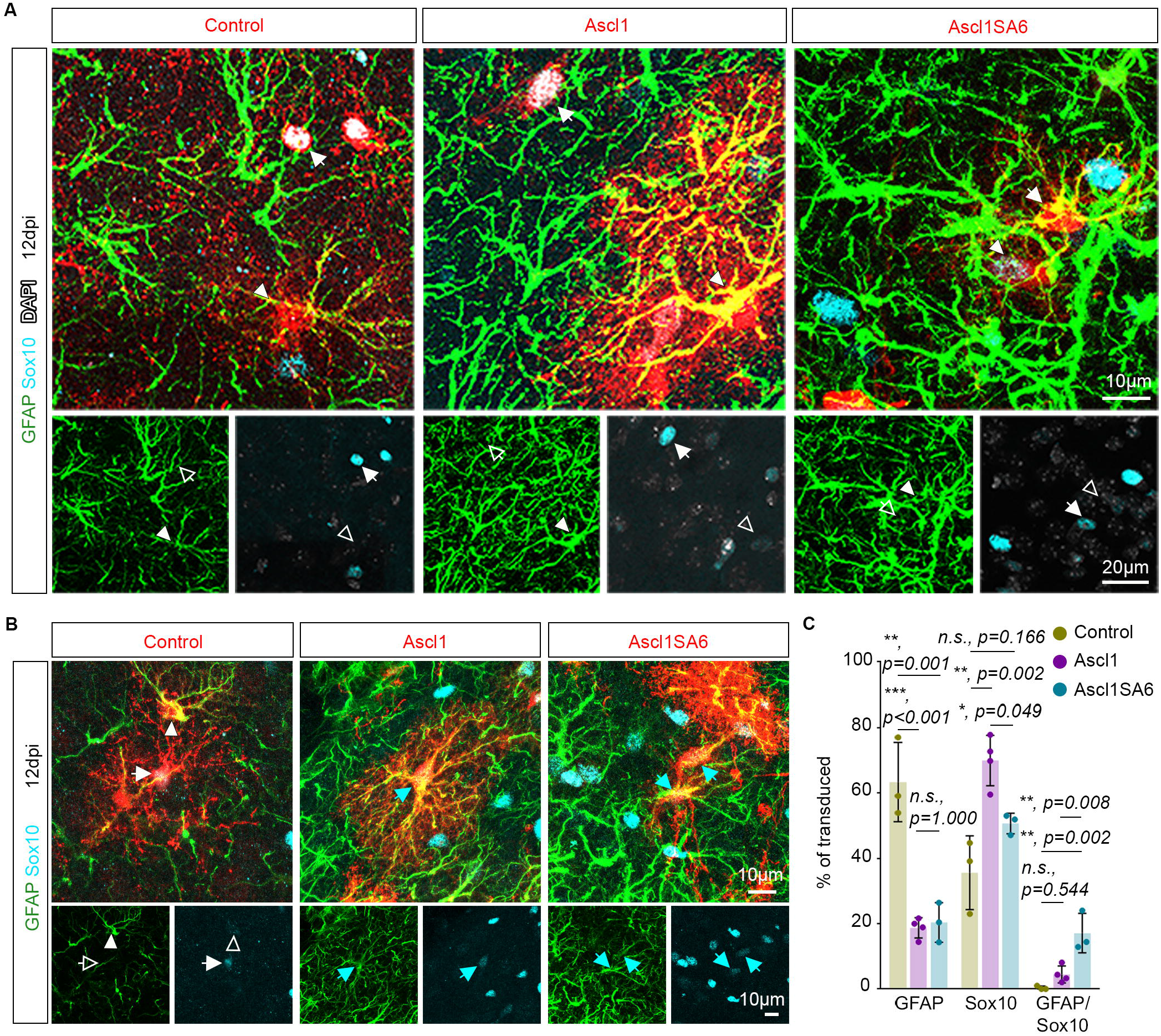
Ascl1 and Ascl1SA6 induce an increase in the number of cells expressing oligodendroglial markers. **(A)** Confocal images depicting control-, Ascl1- and Ascl1SA6-transduced cells (in red) expressing either GFAP (in green, arrowheads) or Sox10 (in cyan, arrows). **(B)** Confocal images depicting Ascl1- and Ascl1SA6-transduced cells (in red) co-expressing (cyan arrows) GFAP (in green) and Sox10 (in cyan). In control, the expression of GFAP (in green, arrowheads) and Sox10 (in cyan, arrows) was exclusive. **(C)** Quantification of the percentage of transduced cells expressing GFAP, Sox10 or both at 12dpi indicates a concomitant reduction in the relative number of cells expressing an astroglial marker and increase in the relative number of cells expressing an oligodendroglial marker upon transduction with either Ascl1 variant. Full arrows/arrowheads indicate marker-positive cells; empty arrows/arrowheads indicate marker-negative cells.

## DISCUSSION

In the present study, we assessed potential reprogramming of glia during their proliferative expansion in the early postnatal cerebral cortex by overexpression of either wild-type Ascl1 or a mutant variant, in which the six conserved serine-proline motifs located outside of the bHLH domain had been mutated (Ascl1SA6), thereby rendering Ascl1 unresponsive to regulation by phosphorylation (Li et al., 2014). We provide evidence that Ascl1SA6 is more efficient than wild-type Ascl1 in converting postnatal astroglia into neurons *in vitro*. Furthermore, we show that the reprogramming efficiency of both Ascl1 and Ascl1SA6 *in vivo* is surprisingly low in the early postnatal cortex compared to the results obtained *in vitro*. Interestingly, while only few Ascl1-transduced cells converted into neurons, we observed a relative shift from GFAP-positive cells to Sox10-positive cells, suggesting an increase in the number of cells of the oligodendroglial lineage at the expense of astroglia.

Our results indicate that Ascl1 reprograms proliferative postnatal cortical glia into neurons with low efficiency *in vivo*. This is in agreement with previous studies reporting inefficient neuronal reprogramming following retrovirus- or lentivirus-mediated expression of Ascl1 in reactive glia in the adult lesioned cortex (Heinrich et al., 2014), adult striatum (Niu et al., 2015) and adult lesioned spinal cord (Su et al., 2014). In contrast to these findings, another study reported very efficient reprogramming of glia into mature neurons following adeno-associated virus (AAV)-mediated expression of Ascl1 in the dorsal midbrain, striatum and somatosensory cortex (Liu et al., 2015). However, misidentification of endogenous neurons as glia-derived neurons was recently reported following AAV-mediated expression of Neurod1, possibly due to transgene sequence-specific effects *in cis* (Wang et al., 2021). Thus, one possible explanation for the apparent discrepancy with regard to the efficiency of Ascl1 to induce reprogramming *in vivo* is that AAV-mediated expression of Ascl1, similarly to Neurod1, resulted in labelling of endogenous neurons. Future studies combining AAV-mediated expression of reprogramming factors such as Ascl1 with genetic lineage tracing are required to clarify the origin of seemingly induced neurons (Wang et al., 2021; Leaman et al., 2022).

The apparent difference in reprogramming potency of Ascl1 *in vitro* and *in vivo* could be attributed to various factors. 1) Enhanced intrinsic cell plasticity of cultured astrocytes as compared to astrocytes *in vivo* despite both being in a similar proliferative state. The protocol employed here to culture and reprogram astrocytes may enhance their competence to undergo cell fate conversion. Indeed, a previous study showed that allowing these astrocytes to mature *in vitro* even only for few days prior to proneural factor activation resulted in a drastic decrease in reprogramming rate, an effect that could be attributed to activation of the REST/coREST repressor complex and accompanying epigenetic maturation (Masserdotti et al., 2015). *In vivo*, REST/coREST complex activity may be already higher, thereby safeguarding glial identity against Ascl1-induced neurogenic reprogramming. 2) Another important difference obviously consists in considerably more complex local environment in which these glial cells find themselves *in vivo*. Nearly nothing is known about the influence that other cell types exert on cells that successfully undergo reprogramming or fail to do so *in vivo*. However, *in vitro* studies have shown that human pericytes undergoing reprogramming by Ascl1 and Sox2 pass through a neural stem cell-like stage during which they become responsive to several intercellular signaling pathways including Notch signaling (Karow et al., 2018). Thus, it is conceivable that signaling molecules as well as extracellular matrix components secreted by cells within the local environment could impinge on early and perhaps more vulnerable reprogramming stages, thereby curtailing progression towards neurogenesis.

Our retroviral vectors were found to target proliferative cells of both the astroglial and oligodendroglial lineage. The overall very low conversion efficiency suggests that not only astroglia, but also cells of the oligodendroglial lineage possess effective safeguarding mechanisms that protect against acquiring a neurogenic fate. In fact, these safeguarding mechanisms are effective even when confronted with a powerful transcription factor with pioneer factor activity, such as Ascl1 (Wapinski et al., 2013; Raposo et al., 2015; Park et al., 2017) or a mutant variant with even increased neurogenic capacity (Woods et al., 2022). Thus, despite being in a proliferative state, astroglia and oligodendrocyte progenitor cells could be potentially less plastic than their adult reactive counterparts in the injured adult brain (Sirko et al., 2013; Magnusson et al., 2014; Faiz et al., 2015; Nato et al., 2015).

While Ascl1 did not induce neurogenic conversion in cells of the astroglial and oligodendroglial lineages, we observed a significant shift in the ratio of virus-transduced astroglial to oligodendroglial cells. Intriguingly, the same shift was observed when using the more neurogenic mutant Ascl1SA6. Several mechanisms could account for that: enhanced expansion of the oligodendroglial cells by activating Ascl1-mediated proliferative programs (Castro et al., 2011); conversely, enhanced cell cycle exit of astrocytes expressing Ascl1; enhanced or reduced survival of oligodendroglial or astroglial lineage cells, respectively; finally, conversion of astroglial cells towards an oligodendroglial fate. The latter would be consistent with the fact that Ascl1 is known to contribute to oligodendrogliogenesis in the developing and adult brain (Parras et al., 2004; Parras et al., 2007). Moreover, studies in the adult hippocampus have previously shown that similar retroviral expression of Ascl1 in neural stem cells, contrary to expectation, promoted oligodendrogliogenesis instead of GABAergic neurogenesis (Jessberger et al., 2008; Braun et al., 2015). Intriguingly, if this latter scenario is indeed the case, the fact that Ascl1SA6 caused a similar shift as wildtype Ascl1 seems to preclude an interpretation according to which acquisition of an oligodendroglial fate would require Ascl1 to be phosphorylated. One line of evidence arguing for at least some oligodendrogliogenic reprogramming by Ascl1 consists in the fact that both variants caused the emergence of GFAP and Sox10 double-positive cells, which were not observed in control virus transduced brains, potentially hinting at an intermediate state between the two glial lineages. Be that as it may, future studies will be needed to distinguish between the mutually non-exclusive mechanisms of action delineated here.

In sum, our study reveals that proliferative glia in the healthy postnatal cerebral cortex are safeguarded against potential neurogenic fate conversion induced by pioneer transcription factors such as Ascl1. Further work will be needed to assess whether additional factors synergizing with Ascl1, such as Sox2 (Heinrich et al., 2014), Dxl2 (Lentini et al., 2021) or Bcl2 (Gascon et al., 2016) could help overcoming these potent safeguarding mechanisms *in vivo*.

## Supporting information

Supplementary Movie 1 (Control)

Supplementary Movie 2 (Ascl1)

Supplementary Movie 3 (Ascl1SA6)

## Author Contributions

Methodology, investigation and formal analysis C.G., N.M., S.P.; Writing – Original Draft, C.G., N.M., S.P.; Funding Acquisition, Conceptualization, Visualization, Writing – Review and Editing, C.S., B.B., S.P. All authors contributed to the article and approved the submitted version.

## Acknowledgments

We are grateful to the members of the Berninger laboratory for their helpful comments and critical feedback over the course of this study. We acknowledge support by the Microscopy Core Facility of the Institute of Molecular Biology (IMB) Mainz.

## Data availability statement

The original contributions presented in the study are included in the article/Material, further inquiries can be directed to the corresponding author.

## REFERENCES

Berninger, B., Costa, M.R., Koch, U., Schroeder, T., Sutor, B., Grothe, B., et al. (2007). Functional properties of neurons derived from in vitro reprogrammed postnatal astroglia. J Neurosci 27(32), 8654–8664. doi: 10.1523/JNEUROSCI.1615-07.2007.

Braun, S.M., Pilz, G.A., Machado, R.A., Moss, J., Becher, B., Toni, N., et al. (2015). Programming Hippocampal Neural Stem/Progenitor Cells into Oligodendrocytes Enhances Remyelination in the Adult Brain after Injury. Cell Rep 11(11), 1679–1685. doi: 10.1016/j.celrep.2015.05.024.

Casarosa, S., Fode, C., and Guillemot, F. (1999). Mash1 regulates neurogenesis in the ventral telencephalon. Development 126(3), 525–534.

Castro, D.S., Martynoga, B., Parras, C., Ramesh, V., Pacary, E., Johnston, C., et al. (2011). A novel function of the proneural factor Ascl1 in progenitor proliferation identified by genome-wide characterization of its targets. Genes Dev 25(9), 930–945. doi: 10.1101/gad.627811.

Chanda, S., Ang, C.E., Davila, J., Pak, C., Mall, M., Lee, Q.Y., et al. (2014). Generation of induced neuronal cells by the single reprogramming factor ASCL1. Stem Cell Reports 3(2), 282–296. doi: 10.1016/j.stemcr.2014.05.020.

Clavreul, S., Abdeladim, L., Hernandez-Garzon, E., Niculescu, D., Durand, J., Ieng, S.H., et al. (2019). Cortical astrocytes develop in a plastic manner at both clonal and cellular levels. Nat Commun 10(1), 4884. doi: 10.1038/s41467-019-12791-5.

Dennis, D.J., Han, S., and Schuurmans, C. (2019). bHLH transcription factors in neural development, disease, and reprogramming. Brain Res 1705, 48–65. doi: 10.1016/j.brainres.2018.03.013.

Denoth-Lippuner, A., and Jessberger, S. (2021). Formation and integration of new neurons in the adult hippocampus. Nat Rev Neurosci 22(4), 223–236. doi: 10.1038/s41583-021-00433-z.

Faiz, M., Sachewsky, N., Gascon, S., Bang, K.W., Morshead, C.M., and Nagy, A. (2015). Adult Neural Stem Cells from the Subventricular Zone Give Rise to Reactive Astrocytes in the Cortex after Stroke. Cell Stem Cell 17(5), 624–634. doi: 10.1016/j.stem.2015.08.002.

Gascon, S., Murenu, E., Masserdotti, G., Ortega, F., Russo, G.L., Petrik, D., et al. (2016). Identification and Successful Negotiation of a Metabolic Checkpoint in Direct Neuronal Reprogramming. Cell Stem Cell 18(3), 396–409. doi: 10.1016/j.stem.2015.12.003.

Ge, W.P., Miyawaki, A., Gage, F.H., Jan, Y.N., and Jan, L.Y. (2012). Local generation of glia is a major astrocyte source in postnatal cortex. Nature 484(7394), 376–380. doi: 10.1038/nature10959.

Guillemot, F., and Hassan, B.A. (2017). Beyond proneural: emerging functions and regulations of proneural proteins. Curr Opin Neurobiol 42, 93–101. doi: 10.1016/j.conb.2016.11.011.

Heinrich, C., Bergami, M., Gascon, S., Lepier, A., Vigano, F., Dimou, L., et al. (2014). Sox2-mediated conversion of NG2 glia into induced neurons in the injured adult cerebral cortex. Stem Cell Reports 3(6), 1000–1014. doi: 10.1016/j.stemcr.2014.10.007.

Heinrich, C., Blum, R., Gascon, S., Masserdotti, G., Tripathi, P., Sanchez, R., et al. (2010). Directing astroglia from the cerebral cortex into subtype specific functional neurons. PLoS Biol 8(5), e1000373. doi: 10.1371/journal.pbio.1000373.

Heinrich, C., Gascon, S., Masserdotti, G., Lepier, A., Sanchez, R., Simon-Ebert, T., et al. (2011). Generation of subtype-specific neurons from postnatal astroglia of the mouse cerebral cortex. Nat Protoc 6(2), 214–228. doi: 10.1038/nprot.2010.188.

Jessberger, S., Toni, N., Clemenson, G.D., Jr., Ray, J., and Gage, F.H. (2008). Directed differentiation of hippocampal stem/progenitor cells in the adult brain. Nat Neurosci 11(8), 888–893. doi: 10.1038/nn.2148.

Karow, M., Camp, J.G., Falk, S., Gerber, T., Pataskar, A., Gac-Santel, M., et al. (2018). Direct pericyte-to-neuron reprogramming via unfolding of a neural stem cell-like program. Nat Neurosci 21(7), 932–940. doi: 10.1038/s41593-018-0168-3.

Karow, M., Sanchez, R., Schichor, C., Masserdotti, G., Ortega, F., Heinrich, C., et al. (2012). Reprogramming of pericyte-derived cells of the adult human brain into induced neuronal cells. Cell Stem Cell 11(4), 471–476. doi: 10.1016/j.stem.2012.07.007.

Leaman, S., Marichal, N., and Berninger, B. (2022). Reprogramming cellular identity in vivo. Development 149(4). doi: 10.1242/dev.200433.

Lentini, C., d’Orange, M., Marichal, N., Trottmann, M.M., Vignoles, R., Foucault, L., et al. (2021). Reprogramming reactive glia into interneurons reduces chronic seizure activity in a mouse model of mesial temporal lobe epilepsy. Cell Stem Cell 28(12), 2104–2121 e2110. doi: 10.1016/j.stem.2021.09.002.

Li, S., Mattar, P., Dixit, R., Lawn, S.O., Wilkinson, G., Kinch, C., et al. (2014). RAS/ERK signaling controls proneural genetic programs in cortical development and gliomagenesis. J Neurosci 34(6), 2169–2190. doi: 10.1523/JNEUROSCI.4077-13.2014.

Liu, Y., Miao, Q., Yuan, J., Han, S., Zhang, P., Li, S., et al. (2015). Ascl1 Converts Dorsal Midbrain Astrocytes into Functional Neurons In Vivo. J Neurosci 35(25), 9336–9355. doi: 10.1523/JNEUROSCI.3975-14.2015.

Magnusson, J.P., Goritz, C., Tatarishvili, J., Dias, D.O., Smith, E.M., Lindvall, O., et al. (2014). A latent neurogenic program in astrocytes regulated by Notch signaling in the mouse. Science 346(6206), 237–241. doi: 10.1126/science.346.6206.237.

Masserdotti, G., Gillotin, S., Sutor, B., Drechsel, D., Irmler, M., Jorgensen, H.F., et al. (2015). Transcriptional Mechanisms of Proneural Factors and REST in Regulating Neuronal Reprogramming of Astrocytes. Cell Stem Cell 17(1), 74–88. doi: 10.1016/j.stem.2015.05.014.

Nato, G., Caramello, A., Trova, S., Avataneo, V., Rolando, C., Taylor, V., et al. (2015). Striatal astrocytes produce neuroblasts in an excitotoxic model of Huntington’s disease. Development 142(5), 840–845. doi: 10.1242/dev.116657.

Niu, W., Zang, T., Smith, D.K., Vue, T.Y., Zou, Y., Bachoo, R., et al. (2015). SOX2 reprograms resident astrocytes into neural progenitors in the adult brain. Stem Cell Reports 4(5), 780–794. doi: 10.1016/j.stemcr.2015.03.006.

Oproescu, A.M., Han, S., and Schuurmans, C. (2021). New Insights Into the Intricacies of Proneural Gene Regulation in the Embryonic and Adult Cerebral Cortex. Front Mol Neurosci 14, 642016. doi: 10.3389/fnmol.2021.642016.

Ory, D.S., Neugeboren, B.A., and Mulligan, R.C. (1996). A stable human-derived packaging cell line for production of high titer retrovirus/vesicular stomatitis virus G pseudotypes. Proc Natl Acad Sci U S A 93(21), 11400–11406. doi: 10.1073/pnas.93.21.11400.

Park, N.I., Guilhamon, P., Desai, K., McAdam, R.F., Langille, E., O’Connor, M., et al. (2017). ASCL1 Reorganizes Chromatin to Direct Neuronal Fate and Suppress Tumorigenicity of Glioblastoma Stem Cells. Cell Stem Cell 21(3), 411. doi: 10.1016/j.stem.2017.08.008.

Parras, C.M., Galli, R., Britz, O., Soares, S., Galichet, C., Battiste, J., et al. (2004). Mash1 specifies neurons and oligodendrocytes in the postnatal brain. EMBO J 23(22), 4495–4505. doi: 10.1038/sj.emboj.7600447.

Parras, C.M., Hunt, C., Sugimori, M., Nakafuku, M., Rowitch, D., and Guillemot, F. (2007). The proneural gene Mash1 specifies an early population of telencephalic oligodendrocytes. J Neurosci 27(16), 4233–4242. doi: 10.1523/JNEUROSCI.0126-07.2007.

Peron, S., and Berninger, B. (2015). Reawakening the sleeping beauty in the adult brain: neurogenesis from parenchymal glia. Curr Opin Genet Dev 34, 46–53. doi: 10.1016/j.gde.2015.07.004.

Poitras, L., Ghanem, N., Hatch, G., and Ekker, M. (2007). The proneural determinant MASH1 regulates forebrain Dlx1/2 expression through the I12b intergenic enhancer. Development 134(9), 1755–1765. doi: 10.1242/dev.02845.

Quan, X.J., Yuan, L., Tiberi, L., Claeys, A., De Geest, N., Yan, J., et al. (2016). Post-translational Control of the Temporal Dynamics of Transcription Factor Activity Regulates Neurogenesis. Cell 164(3), 460–475. doi: 10.1016/j.cell.2015.12.048.

Raposo, A., Vasconcelos, F.F., Drechsel, D., Marie, C., Johnston, C., Dolle, D., et al. (2015). Ascl1 Coordinately Regulates Gene Expression and the Chromatin Landscape during Neurogenesis. Cell Rep 10(9), 1544–1556. doi: 10.1016/j.celrep.2015.02.025.

Sharif, N., Calzolari, F., and Berninger, B. (2021). Direct In Vitro Reprogramming of Astrocytes into Induced Neurons. Methods Mol Biol 2352, 13–29. doi: 10.1007/978-1-0716-1601-7_2.

Sirko, S., Behrendt, G., Johansson, P.A., Tripathi, P., Costa, M., Bek, S., et al. (2013). Reactive glia in the injured brain acquire stem cell properties in response to sonic hedgehog. [corrected]. Cell Stem Cell 12(4), 426–439. doi: 10.1016/j.stem.2013.01.019.

Su, Z., Niu, W., Liu, M.L., Zou, Y., and Zhang, C.L. (2014). In vivo conversion of astrocytes to neurons in the injured adult spinal cord. Nat Commun 5, 3338. doi: 10.1038/ncomms4338.

Wang, L.L., Serrano, C., Zhong, X., Ma, S., Zou, Y., and Zhang, C.L. (2021). Revisiting astrocyte to neuron conversion with lineage tracing in vivo. Cell. doi: 10.1016/j.cell.2021.09.005.

Wapinski, O.L., Vierbuchen, T., Qu, K., Lee, Q.Y., Chanda, S., Fuentes, D.R., et al. (2013). Hierarchical mechanisms for direct reprogramming of fibroblasts to neurons. Cell 155(3), 621–635. doi: 10.1016/j.cell.2013.09.028.

Woods, L.M., Ali, F.R., Gomez, R., Chernukhin, I., Marcos, D., Parkinson, L.M., et al. (2022). Elevated ASCL1 activity creates de novo regulatory elements associated with neuronal differentiation. BMC Genomics 23(1), 255. doi: 10.1186/s12864-022-08495-8.

